# Neuronal programming by microbiota enables environmental regulation of intestinal motility

**DOI:** 10.1101/579250

**Authors:** Yuuki Obata, Stefan Boeing, Álvaro Castaño, Ana Carina Bon-Frauches, Mercedes Gomez de Agüero, Werend Boesmans, Bahtiyar Yilmaz, Rita Lopes, Almaz Huseynova, Muralidhara Rao Maradana, Pieter Vanden Berghe, Andrew J. Murray, Brigitta Stockinger, Andrew J. Macpherson, Vassilis Pachnis

**Author notes:** These authors contributed equally to the work (Stefan Boeing, Álvaro Castaño, Ana Carina Bon-Frauches). Equal contribution (Brigitta Stockinger, Andrew J. Macpherson, Vassilis Pachnis). Lead Contact: Vassilis Pachnis.

## Abstract

Environmental signals modulate the activity of the nervous system and harmonize its output with the outside world. Synaptic activity is crucial for integrating sensory and effector neural pathways but the role of transcriptional mechanisms as environmental sensors in the nervous system remains unclear. By combining a novel strategy for transcriptomic profiling of enteric neurons with microbiota manipulation, we demonstrate that the transcriptional programs of intestinal neural circuits depend on their anatomical and physiological context. We also identify the ligand-dependent transcription factor Aryl hydrocarbon Receptor (AhR) is an intrinsic regulator of enteric nervous system output. AhR is instated as a neuronal biosensor in response to microbiota colonization allowing resident enteric neurons to directly monitor and respond to the intestinal microenvironment. We suggest that AhR signaling integrates neuronal activity with host defence mechanisms towards gut homeostasis and health.

**One Sentence Summary:** Microbiota induce expression of AhR in enteric neurons of the distal intestine enabling them to respond to environmental signals.

---

Integration of gastrointestinal (GI) tissue activity with the dynamic microenvironment of the gut lumen is essential for digestive function and host defence (*1, 2*). The enteric nervous system (ENS) encompasses the intrinsic neural networks of the GI tract regulating many aspects of intestinal physiology, including peristalsis, epithelial cell secretion and mucosal immunity (*2, 3*). In addition to genetic mechanisms and tissue-derived signals (*4*), gut luminal factors, such as diet and microbiota, regulate the development and functional output of the ENS (*5*). Germ-free (GF) mice show morphological changes of intestinal neuronal networks at early postnatal stages (*6*), fail to develop mucosal glial cells (*7*) and display reduced activity of enteric neurons (*8*). Microbiota-diet interactions also influence ENS-dependent peristaltic activity (*9*), while altered composition of microbial communities has been associated with gut motility disorders (*10*). Consistent with these findings, GF and antibiotic-treated mice exhibit reduced gut motility evident by increased transit time (**fig. S1, A and B**) (*11, 12*) and reduced frequency of colonic migrating motor complexes (CMMCs), an ENS-dependent colon-specific motility pattern (**fig. S1C**) (*13, 14*). Interestingly, conventionalization of adult GF (exGF) mice reduces the transit time deficit (**fig. S1A**) (*11*), suggesting that intestinal neural circuits are endowed with molecular mechanisms that continuously monitor the luminal environment of the gut and adjust intestinal motility accordingly. Despite considerable recent progress, the molecular mechanisms whereby the luminal microenvironment regulates ENS physiology and GI homeostasis remain obscure.

We reasoned that genes specifically upregulated in enteric neurons of the colon, the intestinal segment with the highest load of microbiota (*15*), could encode components of signaling cascades activated by luminal signals. To identify such signaling components we first compared the transcriptional profiles of enteric neurons residing within the colon and the small intestine (SI) of specific pathogen-free (SPF) mice. To minimize potential gene expression variation due to diet and other environmental factors, we used mice from two independent animal facilities (Francis Crick Institute, London, UK and University of Bern, Switzerland). Since our pilot studies indicated that the relatively long tissue dissociation protocols commonly used for the purification of intact ENS cells resulted in considerable cell damage and non-specific transcriptional changes, we developed a novel strategy for purification and RNA sequencing of enteric neuron nuclei (nRNAseq). For this, we generated a novel adeno-associated viral (AAV) vector (serotype 9, AAV9) expressing EGFP fused to the KASH nuclear membrane retention domain (EGFPKASH) (*16*) under the control of the neuron-specific CaMKII promoter (**Fig. 1A**). Intravenous administration of AAV9-CaMKII-EGFPKASH particles resulted in nuclear labeling of the majority of enteric neurons within all segments of the intestine (**Fig. 1A and fig. S2A-H**). EGFPKASH-labelled nuclei of myenteric neurons, which are primarily responsible for regulating intestinal motor function, were next isolated from the muscularis externa of the SI and the colon by combining fast and robust protocols of nuclear preparation with fluorescence activated cell sorting (FACS), and subjected to RNAseq (**Fig. 1A and fig. S2I-L**). Bioinformatic analysis demonstrated that the transcriptional profiles of SI and colon myenteric neurons segregated according to their anatomical location and identified sets of genes upregulated in each neuronal population (**Fig. 1B, fig. S2K and table S1**). Among the genes upregulated in colonic neurons were those encoding transcriptional regulators of positional identity (*Hoxd13*), brain development (*Pou3f3*) and environmental sensing (*AhR*), neural tissue-specific proteins (*Col25a1, Syt10*), signaling proteins (*Unc5d, Ano5, Pde1c*) and non-coding RNAs (*Dalir, Pantr2*) (**table S1**). The differential expression of several of these genes in the ENS of the SI and the colon was validated by fluorescence *in situ* hybridization analysis (RNAscope) (**fig. S3**). These experiments demonstrate that discreet transcriptional programs operate in intrinsic neural circuits associated with distinct segments of the mammalian intestine.

**Figure 1.**
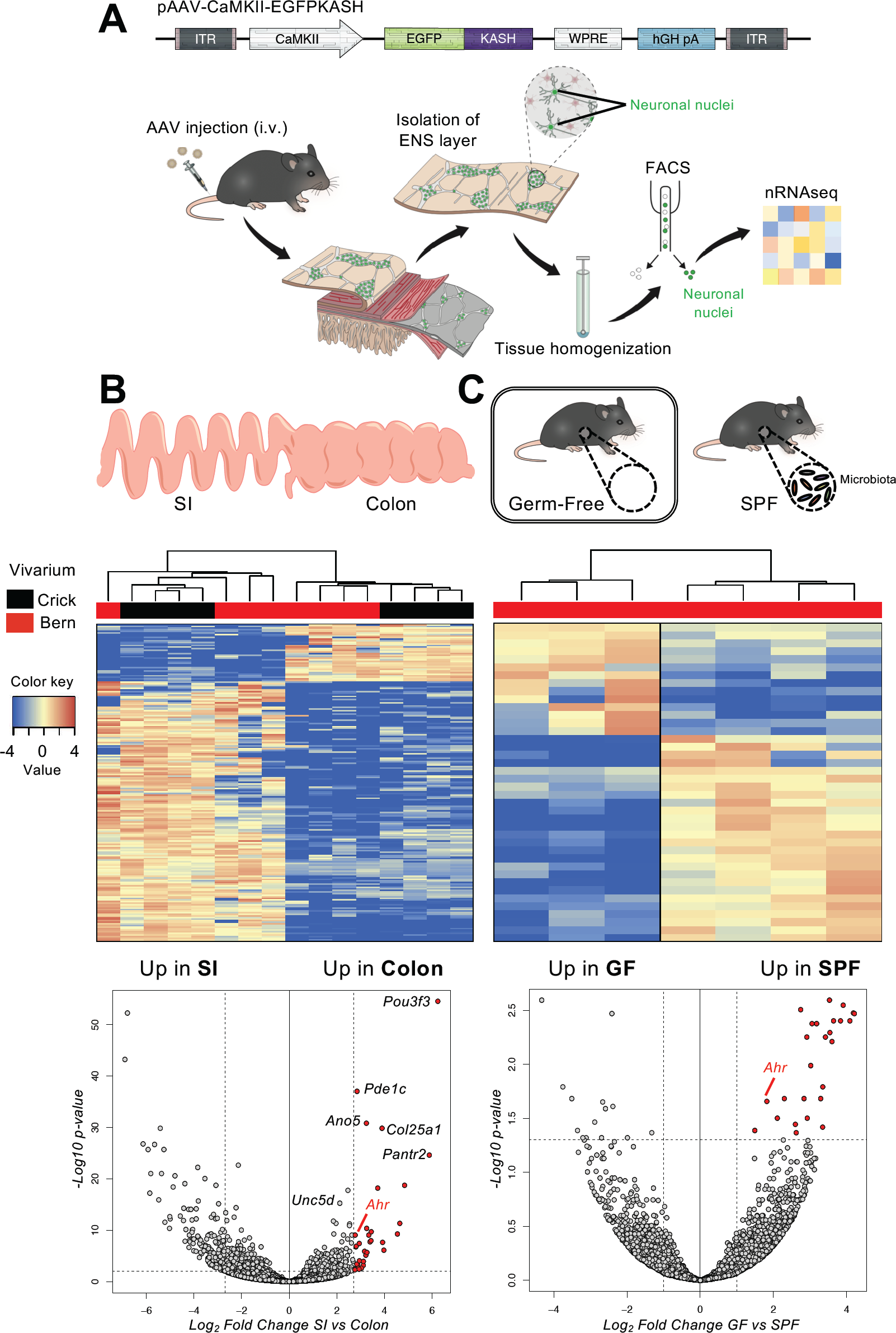
Programming of enteric neuron transcriptome by microbiota. (**A**) Experimental design for AAV-mediated expression of nuclear localized EGFPKASH in enteric neurons and RNAseq of FACS-purified neuronal nuclei. The plasmid used to generate the AAV9-CaMKII-EGFPKASH vector is shown at the top. (**B**) Gene signatures of regional subsets of enteric neurons. Schematic of SI and colon (top). Heat map (middle) shows relative expression levels of selected genes (*logFC ColPos vs SIPos, min < −2.7 and 2.7 < max* with *padj < 0.01;* as indicated in Volcano plot) in 4 Crick and 4 Bern nuclear preparations from SI and colon myenteric neurons. Volcano plot (bottom) shows mean log_2_ fold-change (x axis) and significance (-log_10_ adjusted p-value) of differentially expressed genes between myenteric neurons of the SI and the colon at homeostasis). (**C**) Microbiota-dependent gene signatures of colon myenteric neurons. Neuronal nuclei were isolated from the colon muscularis externa of SPF and GF mice (top). Heat map (middle) shows relative expression levels of select genes (*logFC Colon GF vs Colon SPF min < −1 and 1 > max* with *padj < 0.05;* as indicated in Volcano plot) in myenteric neurons from the colon of SPF and GF mice. Volcano plot (bottom) show mean log_2_ fold-change (x axis) and significance (-log_10_ adjusted p-value) of differentially expressed genes between colon myenteric neurons from SPF and GF mice (bottom).

As an independent experimental strategy for the identification of microbiota-regulated neuronal genes, we used nRNAseq to compare the transcriptional profiles of myenteric neurons from the colon of SPF and GF mice (**Fig 1C**). Despite the presumed effect of microbiota on enteric neuroanatomy (*6, 11*), neuron-specific transcripts were represented equally in the two transcriptomes (**fig. S4A**), indicating that absence of microbiota does not alter the global gene expression profile or organization of intestinal neuronal networks. This idea was further supported by immunostaining for pan-neuronal (HuC/D, PGP9.5, TuJ1 and peripherin) and neuron-subtype (VIP, Calretinin, nNOS, Calbindin, ChAT and NF-M) markers which exhibited similar distribution in the myenteric plexus of SPF and GF mice (**fig. S4, B-F)**. Nevertheless, our transcriptomic analysis identified sets of genes that were specifically upregulated in SPF or GF colonic neurons (**Fig. 1C and table S2**), demonstrating that the microbial communities of the gut modulate the gene expression profile of enteric neurons. Taken together, our studies so far reveal that the anatomical and environmental context of enteric neurons programs their transcriptional output.

One of the genes identified by our nRNAseq experiments as colonic neuron-specific and microbiota-responsive was *AhR* (**Fig. 1, B and C**), suggesting that it is integral to genetic mechanisms employed by enteric neurons to monitor the luminal environment of the colon. *AhR* encodes a member of the bHLH/PAS domain transcription factors, which is activated by a broad range of microbial, dietary and endogenous metabolites (AhR ligands) (*17*). Upon ligand binding, cytosolic AhR translocates into the nucleus and induces expression of, among others, cytochrome P450 (CYP1) enzymes, which metabolize AhR ligands, thereby terminating AhR signaling (*18*). AhR activity in intestinal epithelium and mucosal immune cells is critical for GI physiology and homeostasis (*18, 19*), but the potential role(s) of AhR in enteric neurons is unknown. To investigate the role of AhR in ENS homeostasis and function, we first analyzed its expression profile in enteric neurons. Consistent with our differential gene expression analysis (**Fig. 1B and fig. S3**), virtually all myenteric neurons of the colon identified by pan-neuronal (HuC/D, peripherin) and subtype markers (ChAT, nNOS, Calretinin, Calbindin, or NF-M) were immunostained with AhR-specific antibodies (**Fig. 2, A-E**). Interestingly, the ratio of nuclear versus cytoplasmic signal varied considerably (**Fig 2A**), suggesting asynchronous activation of AhR in enteric neurons. In the muscularis externa of the colon *AhR* was expressed predominantly by myenteric neurons, indicating that neuronal networks in this layer constitute cardinal targets of AhR ligands (**Fig. 2A and fig. S5**). AhR signal was also detected in the myenteric plexus of the distal ileum (which contains the largest number of microorganisms in the SI) and the cecum, but was absent from enteric neurons of the duodenum and jejunum (**Fig. 2F-I**). In contrast to SPF mice, myenteric neurons in the colon of GF and antibiotic-treated animals were labelled weakly with AhR antibodies and had reduced levels of AhR transcripts, but signal was specifically reinstated following colonization of both sets of animals with SPF microbiota (**Fig. 2, J-O and fig. S5A**). Taken together, these studies demonstrate that microbiota colonization regulates *AhR* expression in enteric neurons.

**Figure 2.**
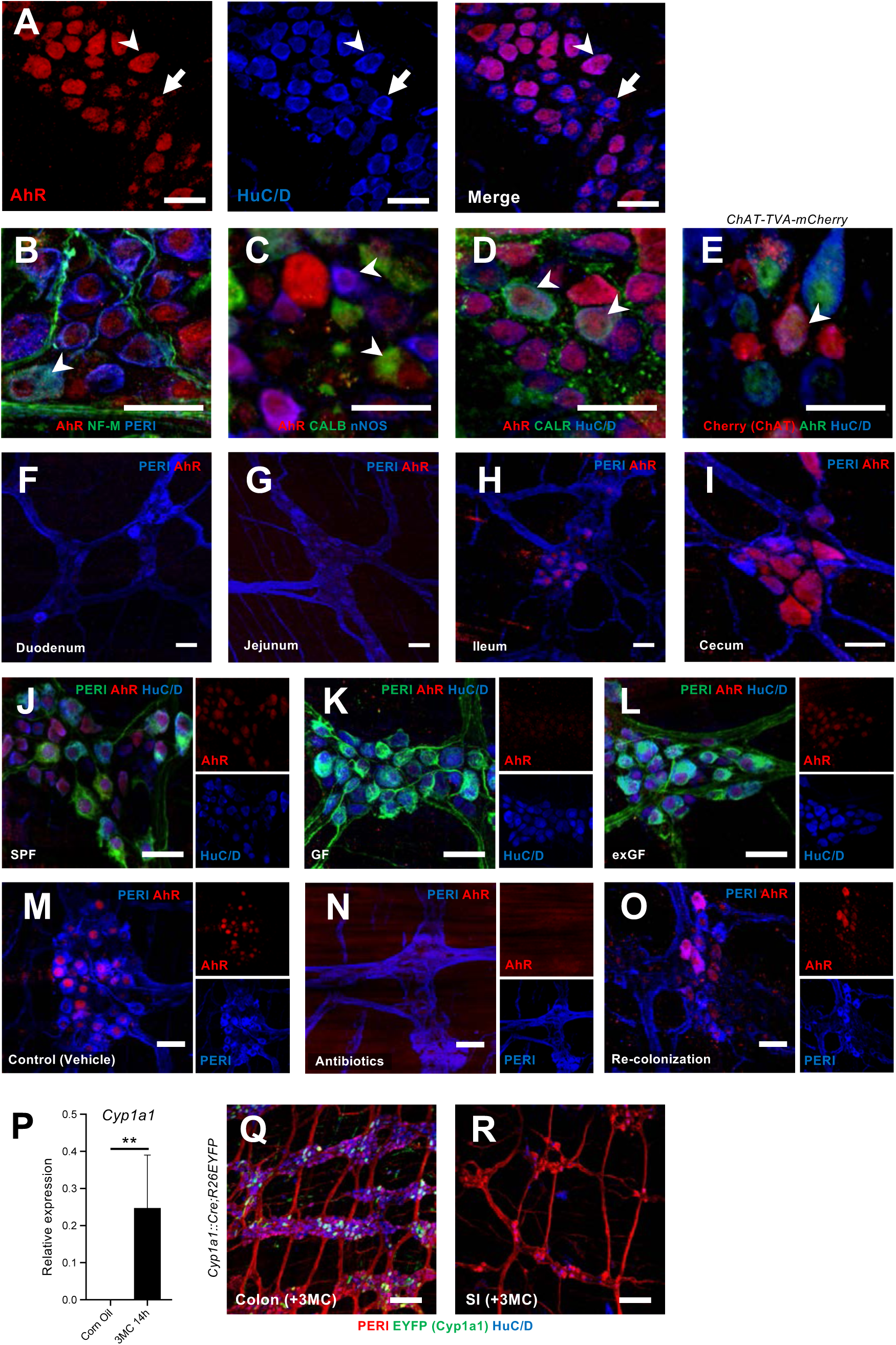
Microbiota-dependent expression and ligand-dependent activation of AhR in myenteric neurons of the colon. (**A**) Immunostaining of colon ganglia for AhR (red) and the pan-neuronal marker HuC/D (blue) in 12 week-old SPF mice (C57Bl/6). AhR signal was either distributed throughout the cell (arrowhead) or was restricted to the nucleus (arrow). Scale bars: 30µm (**B-D**) Immunostaining of neurons from the colon of wild-type (C57Bl/6) mice for AhR (B-D) and the neuronal markers peripherin (PERI) and NF-M (B), calbindin (CALB) and nNOS (C), and calretinin (CALR) and HuC/D (D). AhR signal was detected in all classes of myenteric neurons (arrowheads). Scale bars: 30µm. (**E**) Immunostaining of neurons from the colon of *ChAT-Cherry-TVA* reporter mice for AhR (green), HuC/D (blue) and Cherry (red). Arrowhead indicated an enteric neuron positive for ChAT and AhR. Scale bar: 30µm. (**F-I**) Immunostaining of myenteric ganglia from the duodenum (F), jejunum (G), ileum (H) and cecum (I) with the pan-neuronal marker PERI (blue) and AhR (red). (**J-L**) Immunostaining of myenteric ganglia from the colon of SPF (J), GF (K) and microbiota-colonized GF (exGF; L) mice with the pan-neuronal markers PERI (green) and HuC/D (blue) and AhR (red). Small panels show signal for AhR (top) and HuC/D (bottom). n = 6 animals for each condition. Scale bars: 30µm. (**M-O**) Immunostaining of ganglia from the colon of control (M), antibiotic-treated (N) and microbiota-colonized antibiotic-treated (O) mice with the pan-neuronal marker PERI (blue) and AhR (red). Small panels show signal for AhR (top) and PERI (bottom). n=3 animals for each condition. Scale bars: 30µm. (**P**) qPCR data (means ± SEM) showing levels of Cyp1a1 mRNA in muscularis externa from the colon of control and 3MC-treated wild-type mice. n= 6 control and 8 stimulated animals. Statistical test is the non-parametrical Mann-Whitney U-test \*\**P*<0.01. (**Q** and **R**) Immunostaining of myenteric ganglia from the colon (Q) and SI (R) of 3MC-treated *Cyp1a1::Cre;R26* mice for PERI (red), HuC/D (blue) and EGFP (green). Scale bars: 100µm. 3MC, 3methylchloranthrene.

Earlier reports have suggested that microbiota modulate colonic motility by regulating Toll-like receptor (TLR) signalling (*20*). However, *AhR* expression in colonic neurons is unlikely to depend on TLR signalling, since colonization of GF *Myd88/Trif* double mutant mice (*21*) with microflora induced normal levels of *AhR* expression in the myenteric plexus (**fig. S6A**). Also, T cells, an immune cell subset sensitive to microbial factors, are dispensable for *AhR* expression, since *Tcra*^−/−^ mice (*22*) exhibit normal *AhR* expression in colonic neurons (**fig. S6B**). Therefore, T cells and TLR signaling are dispensable for *AhR* induction in enteric neurons.

To examine whether AhR activity in enteric neurons is subject to ligand regulation, we first compared *Cyp1a1a* transcript levels between control and 3-Methylcholanthrene (3MC)-injected animals (*23*). 14 hours after treatment *Cyp1a1a* mRNA was specifically upregulated in muscularis externa from the colon of animals exposed to 3MC relative to controls (**Fig 2P**). Neuron-specific activation of *AhR* was independently confirmed by GFP immunostaining of muscularis externa preparations from the colon of 3MC-treated *Cyp1a1::Cre;R26EYFP* reporter mice (*18*), in which *Cyp1a1* induction in response to ligand-activated AhR signaling results in permanent expression of EYFP (**Fig 2, Q and R**). Therefore, microbiota-induced expression of *AhR* enables myenteric neurons to respond directly to AhR ligands.

Next, we tested the idea that AhR signaling in enteric neurons is required for regulation of intestinal motility by gut microbiota. For this, we used AAV-mediated gene transfer (**Fig 1A**) to generate mice with enteric neuron-specific deletion of *AhR* (*AhR*^*EN-KO*^). AAV9 vectors expressing Cre recombinase under the control of the CaMKII promoter (AAV9-CaMKII-Cre), were injected intravenously into conditional *AhR* mutant mice (*AhR*^*fl/fl*^) (*24*) carrying the lineage reporter *R26EYFP*, which allowed us to monitor the efficiency and cell type specificity of *AhR* deletion by GFP immunostaining (**Fig. 3A**). Administration of AAV9-CaMKII-Cre to *AhR*^*fl/fl*^*;R26EYFP* mice resulted in EYFP expression and ablation of *AhR* from the majority of enteric neurons (**fig. S7, A and B**). No detectable difference was observed in the number, morphology and neurochemical properties of colonic neurons between control and *AhR*^*EN-KO*^ mice (**fig. S7C**). In addition, constitutive *AhR* mutant mice (*AhR*^*-/-*^) (*25*) showed no deficit in the organization and cellular composition of the ENS (**fig. S7D**). To examine the physiological consequences of *AhR* deletion in colonic neurons, we first compared total intestinal transit time of *AhR*^*EN-KO*^ mice with two sets of control animals: wild-type mice injected with AAV9-CaMKII-Cre (*WT+AAV*) and *AhR*^*fl/fl*^ mice co-housed with the *AhR*^*EN-KO*^ animals to minimize potential microbiota effects. Although *WT+AAV* and *AhR*^*fl/fl*^ mice were indistinguishable in terms of colon histology and motility (**fig. S8, A and B**), *AhR*^*EN-KO*^ animals showed significant increase in gut transit time (**Fig. 3B**), comparable to that observed in microbiota-depleted SPF mice (**fig. S1B**). This phenotype was unlikely to result from ablation of *AhR* in gut-extrinsic sensory neurons, which have been reported to express *AhR* (*26*), since intravenous administration of AAV9-CaMKII vectors showed minimal expression in spinal afferents (**fig. S9A**) and dorsal root ganglia from *AhR*^*EN-KO*^ mice had normal levels of AhR transcripts (**fig. S9B**). To confirm that AhR activity in enteric neurons regulates the physiological output of intestinal neural circuits independently of extrinsic gut innervation, we next used *ex vivo* spatiotemporal mapping of the colon to record neurogenic CMMC activity (*14*). Although CMMCs were indistinguishable between *WT+AAV* and *AhR*^*fl/fl*^ animals (control group), we observed reduced frequency and altered organisation of CMMCs in *AhR*^*EN-KO*^ mice (**Fig. 3C**). We conclude that cell autonomous activity of AhR in enteric neurons regulates the peristaltic activity of the colon.

**Figure 3.**
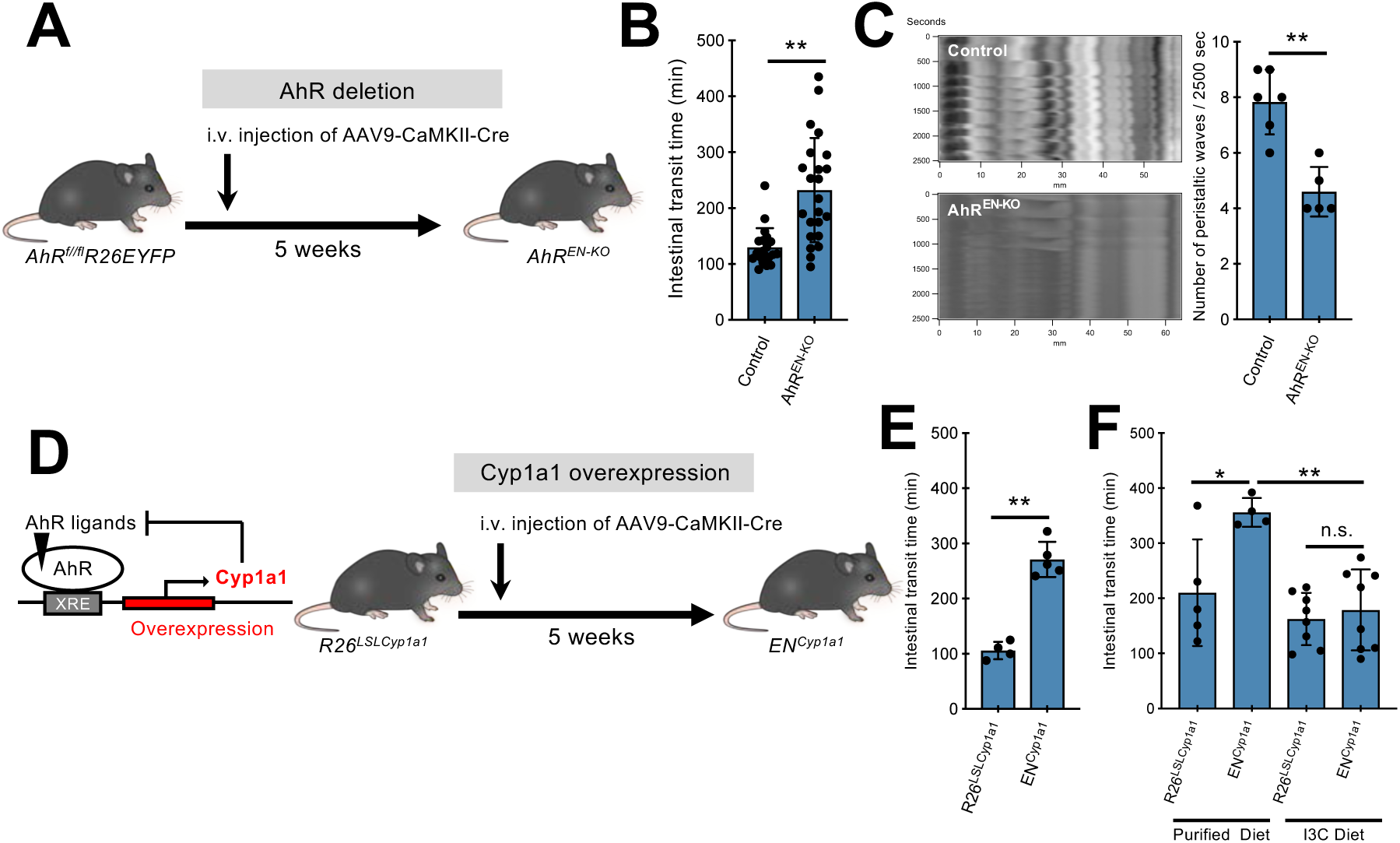
Ligand-induced activation of AhR in enteric neurons regulates intestinal peristalsis. (**A**) Experimental design to delete *AhR* from enteric neurons of *AhR*^*fl/fl*^*;R26EYFP* mice. (**B**) Group data (mean ± SEM) showing the effect of neuron-specific *AhR* deletion on total intestinal transit time. Statistical test is the non-parametrical Mann-Whitney U-test. \*\**P*<0.01. n= 20 (control) and 23 (*AhR*^*EN-KO*^). (**C**) Representative spatiotemporal maps and group data (mean ± SEM) showing the effect of neuron-specific *AhR* deletion on CMMCs recorded *ex vivo*. CMMC, colonic migrating motor complexes. Statistical test is Student’s *t* test. \*\**P*<0.01. n=6 (control) and 5 (*AhR*^*EN-KO*^). (**D**) The negative feedback of CYP1A1 on AhR signaling (left) is the basis for the experimental design (right) to assess the role of *Cyp1a1* overexpression and AhR ligand metabolic clearance in enteric neurons on intestinal motility. (**E**) Group data (mean ± SEM) showing the effect of neuron-specific *Cyp1a1* overexpression on total intestinal transit time. Statistical test is Student’s *t* test. \*\**P*<0.01. n=4 (*R26*^*LSLCyp1a1*^) and 5 (*EN*^*Cyp1a1*^). (**F**) Group data (mean ± SEM) showing that I3C-supplemented diet rescues the total intestinal transit time increase observed in EN^Cyp1a1^ mice. The statistical test was one-way ANOVA followed by Tukey’s test. *\*\* P<0.01. * P<0.05.* n.s., not significant. n=5 (*R26*^*LSLCyp1a1*^ with purified diet), 4 (*EN*^*Cyp1a1*^ with purified diet), 8 (*R26*^*LSLCyp1a1*^ with I3C diet) and 8 (*EN*^*Cyp1a1*^ with I3C diet). I3C, Indole-3-Carbinol.

Metabolic degradation of endogenous AhR ligands by Cyp1a1 is central to the negative feedback regulation of AhR signaling (**Fig. 3D**) (*18*) suggesting that enhanced degradation of AhR ligands by *Cyp1a1* dysregulation would phenocopy the effect of neuron-specific *AhR* deletion on intestinal motility. To test this idea, mice homozygous for the *R26*^*LSL-Cyp1a1*^ allele, which allows conditional overexpression of *Cyp1a1* (*18*), were administered AAV9-CaMKII-Cre vectors resulting in constitutive expression of *Cyp1a1* in enteric neurons (*EN*^*Cyp1a1*^ mice) (**Fig 3D**). Similar to *AhR*^*EN-KO*^ mice, *EN*^*Cyp1a1*^ animals were characterised by increased intestinal transit time (**Fig 3E**) indicating that dysregulation of AhR ligand metabolism in enteric neurons disrupts intestinal neural circuit activity. To examine further the potential role of AhR ligands in neurogenic gut motility, we supplemented the diet of *EN*^*Cyp1a1*^ mice for 4 weeks with the AhR pro-ligand indole-3-carbinol (I3C), which generates the high affinity ligand ICZ (indolo[3,2-b]carbazole) (*27*). Exposure to I3C diet sufficed to rescue the dysmotility in *EN*^*Cyp1a1*^ mice (**Fig 3F**), demonstrating that neuron-specific and ligand-dependent activation of AhR signaling regulates intestinal motility.

To provide direct evidence that AhR signaling is implicated in the regulation of intestinal motility by microbiota (**fig. S1**), antibiotic-treated wild-type mice, which have reduced *AhR* expression in enteric neurons and longer transit time (**Fig 2, M and N, and fig S1B**), were injected with AAV vectors expressing AhR and EGFP under the control of CaMKII (AAV-AhR: AAV9-CaMKII-AhR/EGFP) or control vector (AAV-Ctrl: AAV9-CaMKII-EGFP) and intestinal transit was evaluated 4 weeks later. Since depletion of microbiota was likely to reduce the amount of available AhR ligands (*28*), all animals were fed with I3C-supplemented diet for 1 week prior to the motility assay (**Fig. 4A**). Although antibiotic treatment of mice injected with AAV-Ctrl vector showed a dramatic increase of intestinal transit time, administration of AAV-AhR resulted in significant reduction of total transit time (**Fig 4B**). These experiments demonstrate that enteric neuron-specific AhR signaling is sufficient to reduce intestinal dysmotility secondary to microbiota depletion.

**Figure 4.**
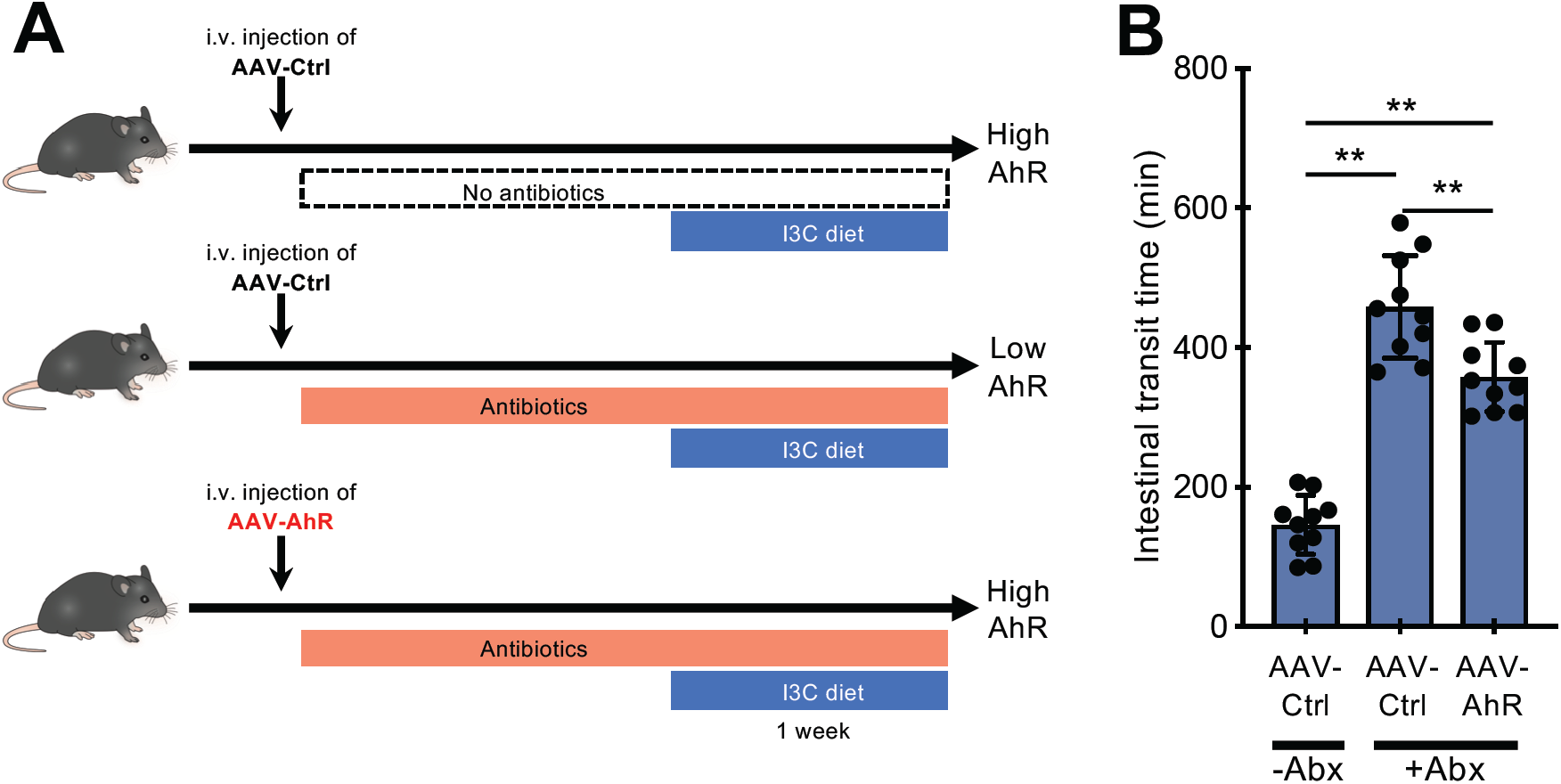
Activation of AhR in enteric neurons promotes intestinal peristalsis in microbiota-depleted mice. (**A**) Experimental design to express and activate AhR in enteric neurons of microbiota-depleted mice. Wild-type SPF mice were injected with control (AAV-Ctrl; top and middle) or AhR-expressing (AAV-AhR; bottom) AAV vectors and treated with vehicle (control; top) or antibiotics (middle and bottom). All animals were fed with I3C-supplemented diet 1 week prior to intestinal transit time analysis. I3C, Indole-3-Carbinol. Abx, Antibiotics (**B**) Group data (mean ± SEM) showing the effect of combinations of AAV-Ctrl and AAV-AhR vectors with antibiotic treatment on total intestinal transit time. Statistical test is one-way ANOVA followed by Tukey’s test. *\*\* p<0.01.* n=10 mice per group.

Digestive physiology and gut homeostasis depend on non-hierarchical input from diverse intestinal tissues (*2*). However, most studies on the homeostatic equilibrium between the gut wall and its lumen have focused on the capacity of intestinal epithelial and immune cells to monitor the microbial and dietary environment and mount appropriate responses to compositional changes and barrier breaching (*29*). Thus, despite the established effect of microbiota and diet on intestinal motility, an important regulator of the luminal microenvironment (*9, 30*), the mechanisms whereby environmental signals regulate the functional output of neural circuits remain largely unexplored. Activation of local sensorimotor reflex units encompassing enterochromaffin cells is a potential mechanism linking microbiota with GI motility (*31*). Here, we describe unique transcriptional programs operating within enteric neurons of distinct intestinal segments, which encode, among others, a novel molecular mechanism that enables neural circuits to directly monitor and respond to changes of the gut microenvironment. This mechanism entails the microbiota-driven installation in enteric neurons of the ligand-dependent transcription factor AhR as a cell autonomous regulatory node, which in turn empowers diverse microbial and non-microbial metabolites to modulate neural output (**fig S10)**. We suggest that AhR is the fulcrum of an ENS-specific surveillance pathway promoting propulsion of luminal contents in response to abnormal microbial overgrowth to prevent dysbiosis and maintain a healthy host-microbe equilibrium (**fig S10**). Therefore, pharmacological activation of AhR in enteric neurons offers a rational therapeutic strategy for the management of bacterial overgrowth resulting from slow gut motility (*32*). In addition to its role in neurogenic motility, AhR-dependent transcriptional programs are central to the barrier function of intestinal epithelial cells and the mucosal immune system. A common signaling pathway operating within diverse cell types offers an effective mechanism to integrate the activity of functionally distinct intestinal tissues towards gut homeostasis and host defence. Perhaps, in a similar manner, metabolite-induced AhR activation in distinct cell lineages of the central nervous system, including neurons (*26, 33*) and microglia (*34*) could integrate their activity under physiological conditions or their response to tissue pathology. In summary, our studies provide mechanistic insight into the environmental adaptation of neural circuits and its role in nervous system homeostasis.

## Acknowledgements

We thank the Crick Science Technology Platforms for expert support with experiments. We also thank Reena Lasrado and Song Hui Chng with tissue dissection and all members of the Pachnis Lab. for insightful comments on the manuscript. Y.O. has been supported by an EMBO long-term fellowship (ALTF 1214-2015), an HFSP postdoctoral fellowship (LT000176/2016) and a travel grant from Boehringer Ingelheim Fonds. This work was supported by the Medical Research Council (MRC) and The Francis Crick Institute (which receives funding from the MRC, Cancer Research UK, and the Wellcome Trust). V.P. was also funded by BBSRC (BB/L022974) and the Wellcome Trust (212300/Z/18/Z).

## SUPPLEMENTARY MATERIALS

## Materials and methods

### Animals

All animal procedures at the Francis Crick Institute were carried out in accordance with the regulatory standards of the UK Home Office and approved by the local Animal Welfare and Ethics Review Body (AWERB). Procedures at the University of Bern were performed in accordance with Swiss Federal Regulations.

The following transgenic lines have been described previously: *Cyp1a1::Cre (35), AhR*^*-/-*^ *(25), AhR*^*fl/fl*^ *(24), R26*^*LSL-Cyp1a1*^ (*18*), *Tcrα*^*-/-*^ (*22*) and *R26EYFP (36)*. Details about the generation of the *ChAT-TVA-mCherry* will be reported elsewhere. Information about this line is available upon request. Transgenic and wild-type *C57BL/6* mice were bred and maintained in the SPF facility of the Francis Crick Institute. For most experiments, animals were on standard Crick diet. For the motility rescue experiment (Fig. 3F), animals were fed with either purified diet (AIN; sniff spezialidäten, GmbH) or purified diet supplemented with Indole-3-Carbinol (I3C, 200 mg/kg; sniff spezialidäten, GmbH).

Wild-type and *Myd88/Trif* double mutant (*21*) GF mice (C57BL/6) were bred and maintained in flexible-film isolators at the Clean Mouse Facility of the University of Bern (Switzerland). GF status was monitored routinely by culture based and other methods and all mice were independently confirmed to be microbe-free (*37*). For bacterial colonisation experiments, fecal contents of SPF mice were orally administered to wild-type and *Myd88/Trif* double mutant GF mice and colonised animals were co-housed and maintained in the SPF facility of the University of Bern for 4 weeks prior the analysis.

For depletion of microbiota, wild-type (C57BL/6) mice were administered a 4-mM acetic acid solution containing 1 g/L ampicillin sodium, 0.5 g/L vancomycin hydrochloride, 1 g/L neomycin sulfate, 1 g/L metronidazole (all from Sigma-Aldrich) and 1% (v/w) artificial sweet flavour (Vimto) via drinking water for 18 days (**Fig. 4**) or 30 days (**fig. S1B**).

For systemic administration of AAV (*38*), 6-week-old mice were intravenously injected with AAV particles (dPCR-GC titre >1e12 GC) in 5% sucrose buffer (150µL per animal).

### Generation of AAV vectors

For the generation of AAV9-CaMKII-EGFPKASH vector, the DNA cassette encoding KASH-tagged EGFP (EGFPKASH) (*16*) was amplified from the PX552 plasmid (Addgene #60958) by Phusion High-Fidelity DNA polymerase. PCR was performed using the 5’-GCTATCGGATCCGCCACCATGGTG-3’ and 5’-GCTATCGAATTCCTAGGTGGGAGG-3’ primers, which allowed us to use BamHI and EcoRI restriction site to substitute EGFPKASH for EGFP in plasmid pAAV-CaMKII-EGFP (Addgene#50469), thus generating pAAV-CaMKII-EGFPKASH (schematic in **Fig. 1A**). NEB-Stable Competent *E. coli* (C3040I) were transformed with pAAV-CaMKII-EGFPKASH and bacteria were grown on LB agar plates containing ampicillin (50 mg/mL). The presence of inverted terminal repeats (ITR) was confirmed by SmaI digestion. Production of AAV particles was carried out as previously described (*39*). Briefly, 293AAV cells were transfected with three plasmids, pAAV-CaMKII-EGFPKASH, pFdelta6 (Adenovirus-helper plasmid) and the capsid plasmid (including the AAV9 Rep and Cap sequences), in Iscove’s modified Dulbecco’s medium (IMDM) containing 5% fetal calf serum using TurboFect transfection reagents (Thermo Fisher Scientific). 16 hours after transfection, IMDM was replaced with Dulbecco’s modified Eagle medium (DMEM). 72 hours after transfection, cells were washed with warm PBS, gently removed from the plate and collected in 50 mL tubes. Lysing of cells and particle purification were carried out using AAVpro® Purification Kit (All Serotypes, TAKARA) according to the manufacturer’s instructions. The purity of the viral particles was assessed by SDS-PAGE after concentration and sterile filtration of viral particles. pAAV-CaMKII-EGFPKASH will be deposited in Addgene.

Large-scale packaging and generation of AAV9-CaMKII-EGFPKASH, AAV9-CaMKII-Cre (Addgene#105558) and AAV9-CaMKII-EGFP (Addgene #105541) vectors was carried out by Penn Vector Core (University of Pennsylvania, USA). AAV9-CaMKII-AhR/EGFP was generated by Vector Biolabs (PA, USA).

### Histopathological analysis

Intestinal segments were harvested from mice infected with AAV9 vectors, fixed with 4% PFA/PBS overnight (O/N) at 4°C and embedded in paraffin. Paraffin was removed from sections with xylene, rehydrated with ethanol and stained with either hematoxylin-eosin (H&E) or Alcian blue-PAS hematoxylin.

### Immunohistochemistry

For immunostaining, the intestine was flushed free of luminal contents, cut along the mesenteric border following removal of adipose tissues on serosa and fixed with 4% paraformaldehyde (PFA) O/N at 4°C. For the preparation of submucosal plexus (SMP) layer, a 1-mL pipette was inserted into the lumen to fully extend the smooth muscle layer containing myenteric plexus (MP), which was removed from mucosal compartment using cotton buds as previously described (*40*). The SMP layer was treated with 30mM EDTA/PBS for 30 min on ice to remove epithelial layer, stretched on Sylgard coated petri dish, and fixed with 4% PFA for 3 hours at 4°C. The fixed tissues were then rinsed with PBS for three times at room temperature. Gut tissues were then permeabilized and pre-blocked with 10% normal donkey serum (NDS)/1% Triton-X100 for 1 hour at room temperature and incubated with primary antibodies (**table S3**) in 10% NDS/1% Triton-X100 for 48 hours (at 4°C). Tissue was then incubated with secondary antibodies (**table S3**) in 10% NDS/1% Triton-X100 for 12 hours at room temperature. DAPI (Molecular Probes #d3571) was used for counter staining. Samples were washed with PBS before mounting with VECTASHIELD (Vector Laboratories).

### Fluorescence in situ hybridization

Fluorescence in situ hybridization on the myenteric plexus was carried out using the Advanced Cell Diagnostics RNAscope® Fluorescent Multiplex Kit (ACD #320850) according to manufacturer’s specification. Briefly, the muscularis externa layer of the gut was dehydrated by serial ethanol treatments and treated with RNAscope® Protease III for 30 min (for SI) or 45 min (for colon) at RT. Tissue was then incubated O/N (at 40°C) with fluorescent probes, 3-Plex Positive Control Probe, 3-Plex Negative Control Probe or customized probes (listed in **table S3**). Following hybridization, tissue was washed twice with Wash Buffer and then subjected to sequential hybridization with pre-amplifier, amplifier DNA (Amp1-FL, Amp 2-FL and Amp 3-FL) and fluorophore (Amp 4 Alt A-FL) at 40°C for 15-30 min for each step. After hybridization, tissues were counterstained with RNAscope® (ACD #320858) and mounted on Superfrost Plus Adhesion Microscope Slides (ThermoFisher Scientific #10149870) using VectaMount Permanent Mounting Medium (ACD #321584).

### Image processing

Immunostained gut preparations were examined with Olympus FV3000-Invert (SW312-CB1) confocal laser scanning microscope and FV31S-SW software (Olympus) using standard excitation and emission filters for visualizing DAPI, Alexa Fluor 488, Alexa Fluor 568, and Alexa Fluor 647. All images were processed with Adobe Photoshop CS 8.0 (Adobe Systems) while analyses were performed using the image-processing package Fiji/ImageJ (Wayne Rasband, NIH).

### Q-PCR

Total RNA was isolated from colonic muscular layer using Trizol LS reagent and the PureLink RNA Micro Kit (Invitrogen #12183016) according to the manufacturer’s specifications and was subjected to reverse transcription using the High-Capacity cDNA Reverse Transcription Kit (Applied Biosystems #4368814). Q-PCR was performed with complementary DNA (cDNA) using Taqman fast universal 2 × PCR Master Mix (Applied Biosystems) and Taqman probes (Applied Biosystems) for *β-actin* (Mm02619580_g1) and *Cyp1a1* (Mm00487218_m1). *Ct* values obtained were normalised to *β-actin*.

### Purification of neuronal nuclei from myenteric plexus

5 weeks after intravenous injection of *AAV9-CaMKII-EGFPKASH*, the longitudinal smooth muscle layer and associated myenteric plexus were peeled off the wall of the SI and colon and subjected to Dounce homogenization in lysis buffer (250mM sucrose, 25mM KCl, 5mM MgCl_2_, 10mM Tris buffer with pH8.0, 1µM DTT) containing 0.1% Triton-X, cOmplete™ EDTA-free protease inhibitor (Sigma-Aldrich) and DAPI. After filtering the homogenate to remove large debris, samples were centrifuged at 1000 × G for 10 min at 4°C to obtain a pellet containing muscularis externa nuclei. For flow cytometric analysis, doublet discrimination gating was applied to exclude aggregated nuclei, and intact nuclei were determined by subsequent gating on the area and height of DAPI intensity. Both EGFP^+^ and EGFP^−^ nuclear populations were collected directly into 1.5mL tube containing Trizol LS reagent (Invitrogen) using an Aria Fusion cell sorter (BD Biosciences). The obtained FCS data were further analyzed using FlowJo software version 10.5.3.

### RNA-sequencing and Bioinformatic analysis

Nuclei-derived RNA was isolated using PureLink RNA Micro Kit (Invitrogen #12183016) according to the manufacturer’s instructions. Double stranded full-length cDNA was generated using the Ovation RNA-Seq System V2 (NuGen Technologies, Inc.). Following quantification on a Qubit 3.0 fluorometer (Thermo Fisher Scientific, Inc.), cDNA was fragmented to 200bp by acoustic shearing using Covaris E220 instrument (Covaris, Inc.) at standard settings. The fragmented cDNA was then normalized to 100ng, which was used for sequencing library preparation using the Ovation Ultralow System V2 1-96 protocol (NuGen Technologies, Inc.). A total of 7 PCR cycles were used for library amplification. The quality and quantity of the final libraries were assessed with TapeStation D1000 Assay (Agilent Technologies, Inc.). The libraries were then normalized to 2,5 nM, pooled and loaded onto a HiSeq4000 (Illumina, Inc.) to generate 75bp single-end reads.

For the bioinformatics analysis, the ‘Trim Galore!’ utility version 0.4.2 was used to remove sequencing adaptors and to quality trim individual reads with the q-parameter set to 20. The sequencing reads were then aligned to the mouse genome and transcriptome (Ensembl GRCm38 release-86) using RSEM version 1.3.0 in conjunction with the STAR aligner version 2.5.2. Sequencing quality of individual samples was assessed using FASTQC version 0.11.5 and RNA-SeQC version 1.1.8. Differential gene expression was determined using the R-Bioconductor package DESeq2 version 1.14.1.

### Intestinal transit time assay

The total intestinal transit time was measured as previously described (*40*). Animals were placed individually in bedding-free cages with the diet and HydroGel (Clear H_2_O). 250µL of 6% (w/v) carmine red dye (Sigma-Aldrich) in 0.5% (w/v) methylcellulose (Sigma-Aldrich) was orally administered to each mouse at 9:00 am in all experiments. The time period from gavage till the emergence of the first red-colored pellet was recorded as total transit time.

### Live video imaging and spatiotemporal mapping of colonic motility

*Ex vivo* video imaging and analysis of colonic motility was performed as described previously (*40*). Briefly, the entire colon was carefully isolated and loosely pinned in an organ bath chamber continuously superfused (flow rate: 4 mL per minute) with oxygenated (95% O2 and 5% CO2) Krebs solution (in mM: 120.9 NaCl, 5.9 KCl, 1.2 MgCl2, 2.5 CaCl2, 1.2 NaH2PO4, 14.4 NaHCO3, 11.5 Glucose) kept at 37°C. Following equilibration of the colon for 30 min, movies of colonic motility were captured (2.5 Hz frame rate) with a QICAM-Fast camera using QCapture Pro 6.0 software (Q-Imaging). Images were read into an Igor Pro (WaveMetrics) and analysed using custom-written algorithms. The bowel’s edges were determined and the width computed and mapped over time. From the generated spatiotemporal maps, the frequency of propagating contractions was determined.

### Statistical analysis

Statistical comparisons between samples were performed in GraphPad Prism software using Student’s t test or one-way ANOVA. A *p* value less than 0.05 was considered statistically significant. No samples were excluded from the statistical analyses.

## Supplemental figure legends

**Figure S1.**
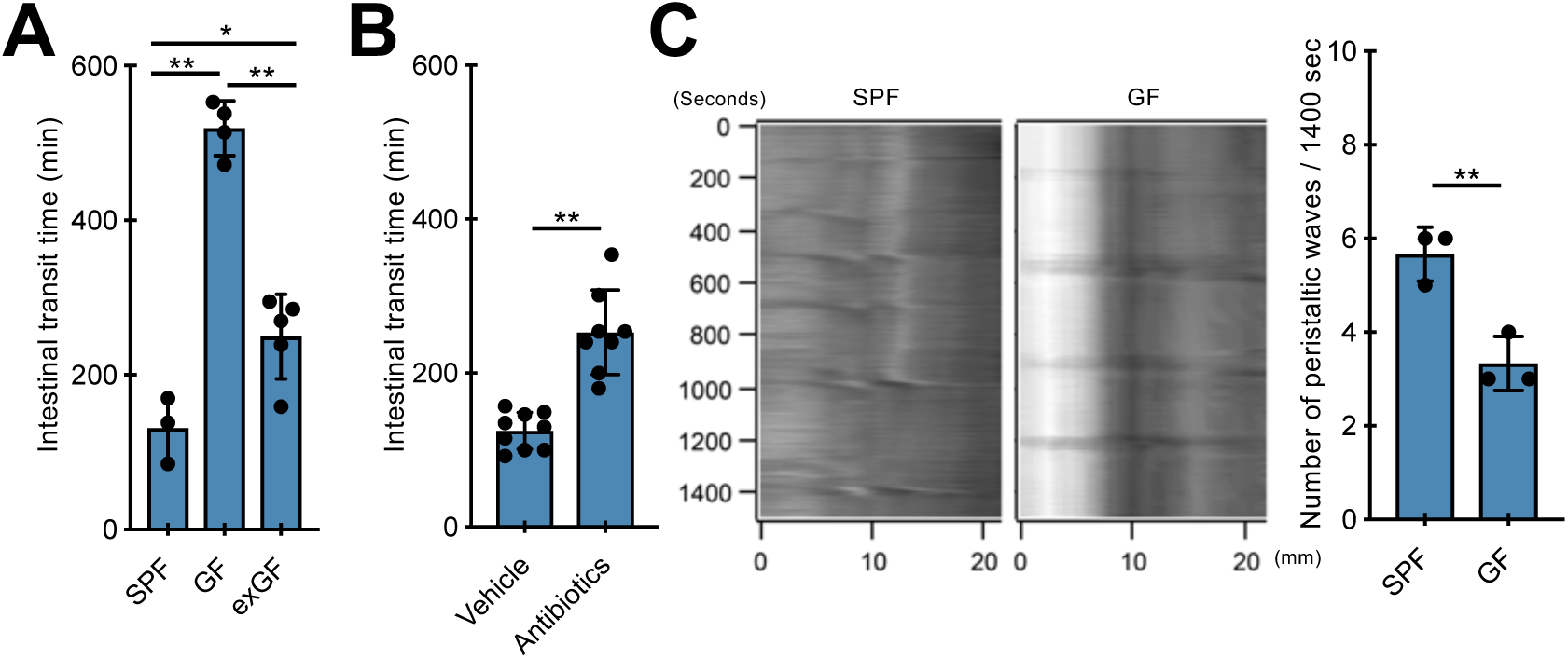
Impaired intestinal motility of microbiota-depleted mice. (**A**) Group data (mean ± SEM) comparing total intestinal transit time between SPF, GF and exGF (GF mice colonized with SPF microbiota) mice. *p* values determined by one-way ANOVA test followed by Tukey’s test. *\*\* p<0.01. * p<0.05.* n=3 (SPF), 4 (GF) and 5 (exGF). (**B**) Group data (mean ± SEM) showing the effect of antibiotic treatment on total intestinal transit time. Statistical test is the non-parametrical Mann-Whitney U-test \*\**P*<0.01. n=9 (vehicle) and 8 (Antibiotics). (**C**) Representative spatiotemporal maps (left) and group data (mean ± SEM; right) showing the effect of microbiota depletion on CMMCs recorded *ex vivo*. Statistical test is Student’s *t* test. \*\**P*<0.01. n=3 mice per group.

**Figure S2.**
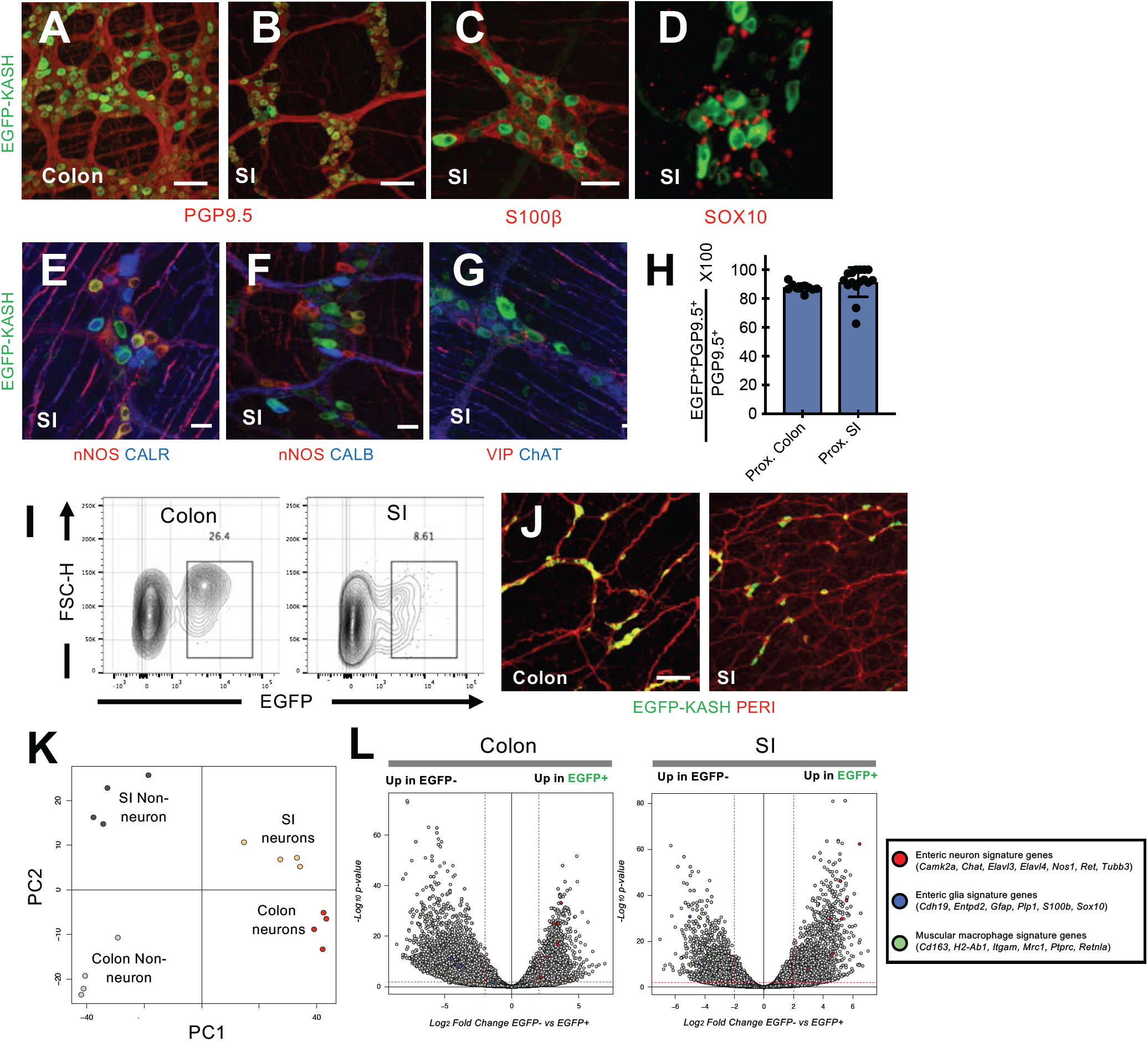
AAV-based transcriptional profiling of enteric neurons. **(A-G)** Representative images of enteric ganglia labelled by intravenous injection of AAV9-CaMKII-EGFPKASH vectors. Colon (A) and SI (B-G) myenteric plexus preparations were immunostained with antibodies against GFP (A-G), PGP9.5 (A and B), S100β (C), SOX10 (D), nNOS and CALR (E), nNOS and CALB (F) and VIP and ChAT (G). Data are representative of two independent experiments. Scale Bars: 100µm (A-D) and 30µm (E-G). CALR, Calretinin; CALB, Calbindin. (**H**) Group data (mean ± SEM) showing that following intravenous administration of the AAV9-CaMKII-EGFPKASH vector the vast majority of PGP9.5^+^ enteric neurons in the proximal SI and colon express EYFP. (**I**) Representative FACS plots of muscularis externa nuclei from the colon (left) and SI (right) of mice injected systemically with AAV9-CaMKII-EGFPKASH. Representative FACS plots gated on single intact DAPI^+^ nuclei. (**J**) Peripherin (PERI; red) and GFP (green) immunostaining of the submucosal plexus remaining after isolation dissection of the muscularis externa (for nRNAseq) from the colon (left) and SI (right) of AAV9-CaMKII-EGFPKASH-injected mice. Images are representative of two independent experiments. Scale Bars: 100µm. (**K**) Principal component analysis of the transcriptomes of EGFP^+^ and EGFP^−^ nuclei isolated from the muscularis externa of the colon and SI of mice injected with AAV9-CaMKII-EGFPKASH vectors. Segregation of transcriptomes according to the neuronal vs non-neuronal origin of the nuclei and anatomical location along the gut. (**L**) Volcano plot showing mean log_2_ fold-change (x axis) and significance (−log_10_ adjusted p-value) of differentially expressed genes between EGFP^+^ and EGFP^−^ nuclei isolated from the colon (left) and SI (right) of mice injected with AAV9-CaMKII-EGFPKASH vectors. Red, blue and green dots indicate the genes specific to enteric neurons (*Ret, Chat, Camk2, Elav3, Nos1, Tubb3*), glial cells (*Sox10, Gfap, Cdh19, Entpd2, Plp1*) (*41*) and muscular macrophages (*CD11b, CD163, MHC class II, Mrc1, Retnla*) (*42*), respectively.

**Figure S3.**
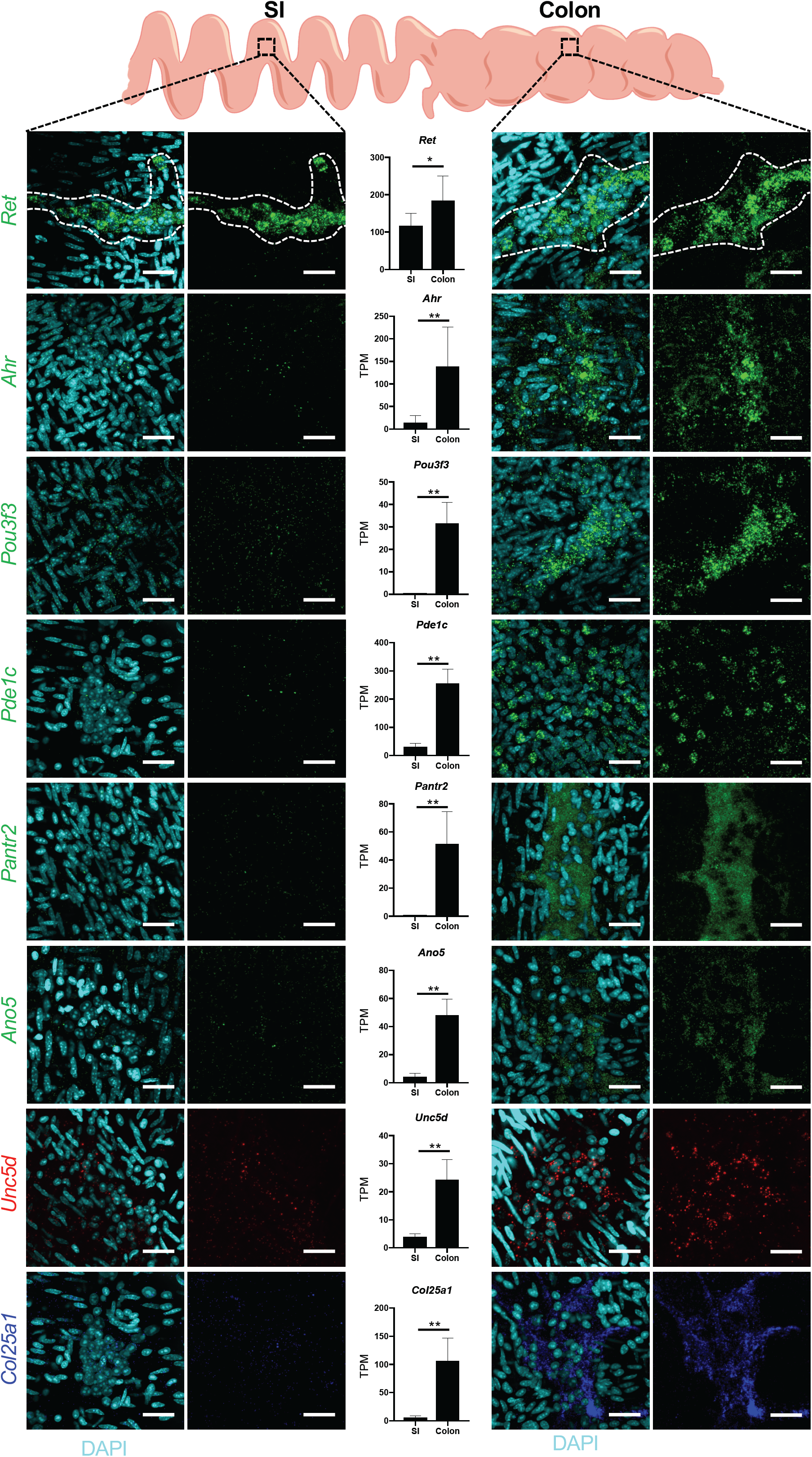
Differential expression of enteric neuron-specific genes in muscularis externa preparations from the SI and colon. Representative images of muscularis externa from SI (left) and colon (right) hybridized with the indicated RNAscope probes and counterstained for DAPI. With the exception of *Ret*, all other genes analyzed here (*Ahr, Pou3f3, Pde1c, Pantr2, Ano5, Unc5d, Col25a1*) were identified by nRNAseq (Fig. 1B) as colon neuron-specific. Quantification of expression levels for each gene in the nRNAseq experiments (TPM values) is also shown. *Ret* is expressed in neurons of myenteric ganglia (outlined by dotted line) of both the SI and the colon (although our quantification shows higher representation in colon samples). *Ahr, Pou3f3, Pde1c, Pantr2, Ano5, Unc5d, Col25a1* are expressed predominantly by myenteric neurons of the colon. Statistical test was Student’s *t* test. When variances were not homogeneous, the data were analyzed by the nonparametrical Mann-Whitney U-test \*\**P*<0.01. TPM, transcripts per kilobase million.

**Figure S4.**
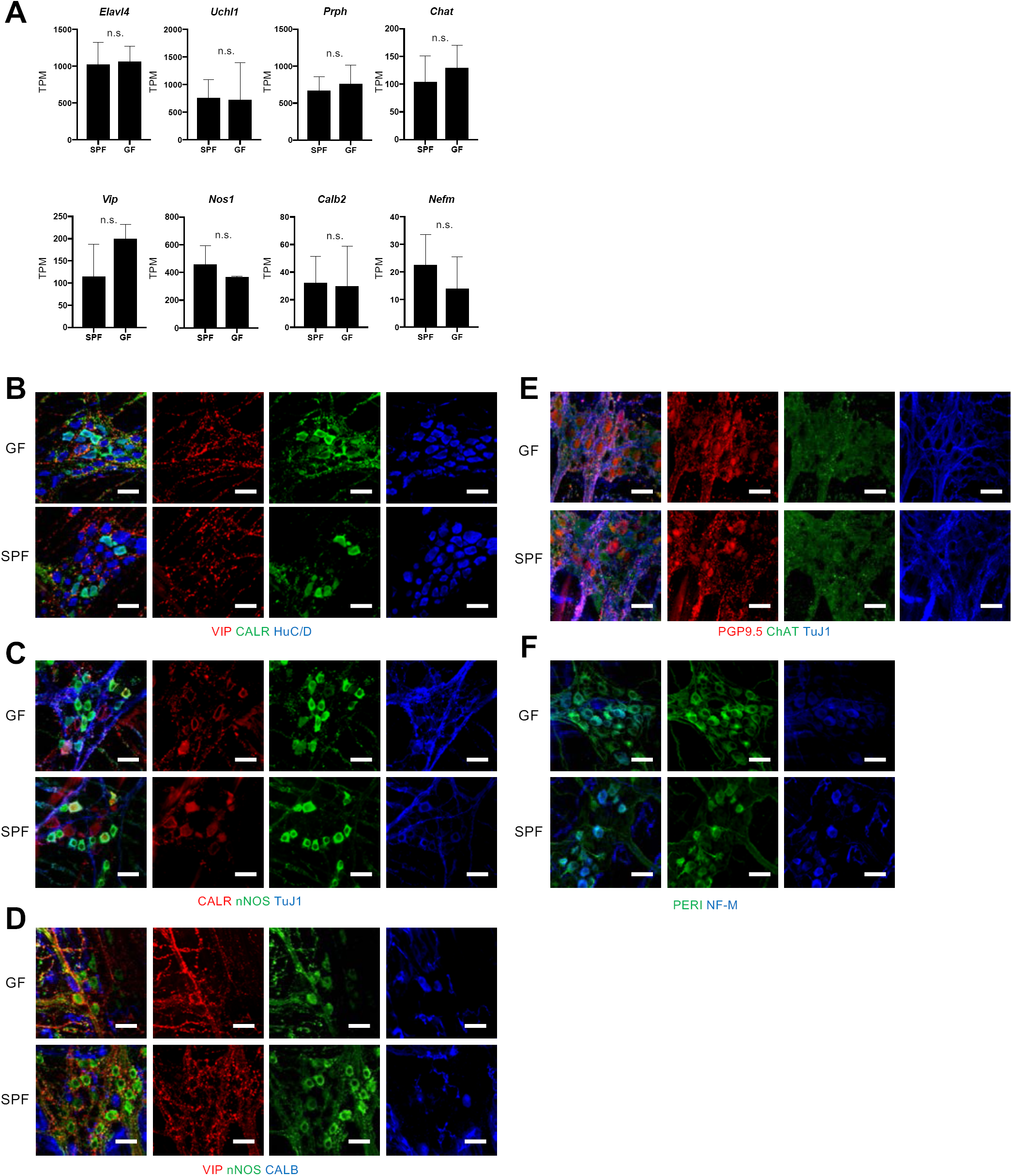
Molecular and neurochemical characterization of colonic neurons in GF and SPF mice. (**A**) Quantification of expression levels (TPM mean ± SEM, n=4 SPF and 3 GF mice) of neuronal gene markers *Elavl4, Uchl1, Prph, Chat, Vip, Nos1, Calb2, Nefm* in the muscularis externa of the colon of SPF and GF mice. Statistical test is Student’s *t* test. When variances were not homogeneous, the data were analyzed by the nonparametrical Mann-Whitney U-test. n.s: not significant. TPM, transcripts per kilobase million. (**B-F**) Immunostaining of colonic myenteric ganglia from SPF and GF mice with VIP, CALR and HuC/D (B), CALR, nNOS and TuJ1 (C), VIP, nNOS and CALB (D), PGP9.5, ChAT and TuJ1 (E) and PERI and NF-M (F). Scale Bars: 30µm. CALR, Calretinin; CALB, Calbindin; PERI, peripherin.

**Figure S5.**
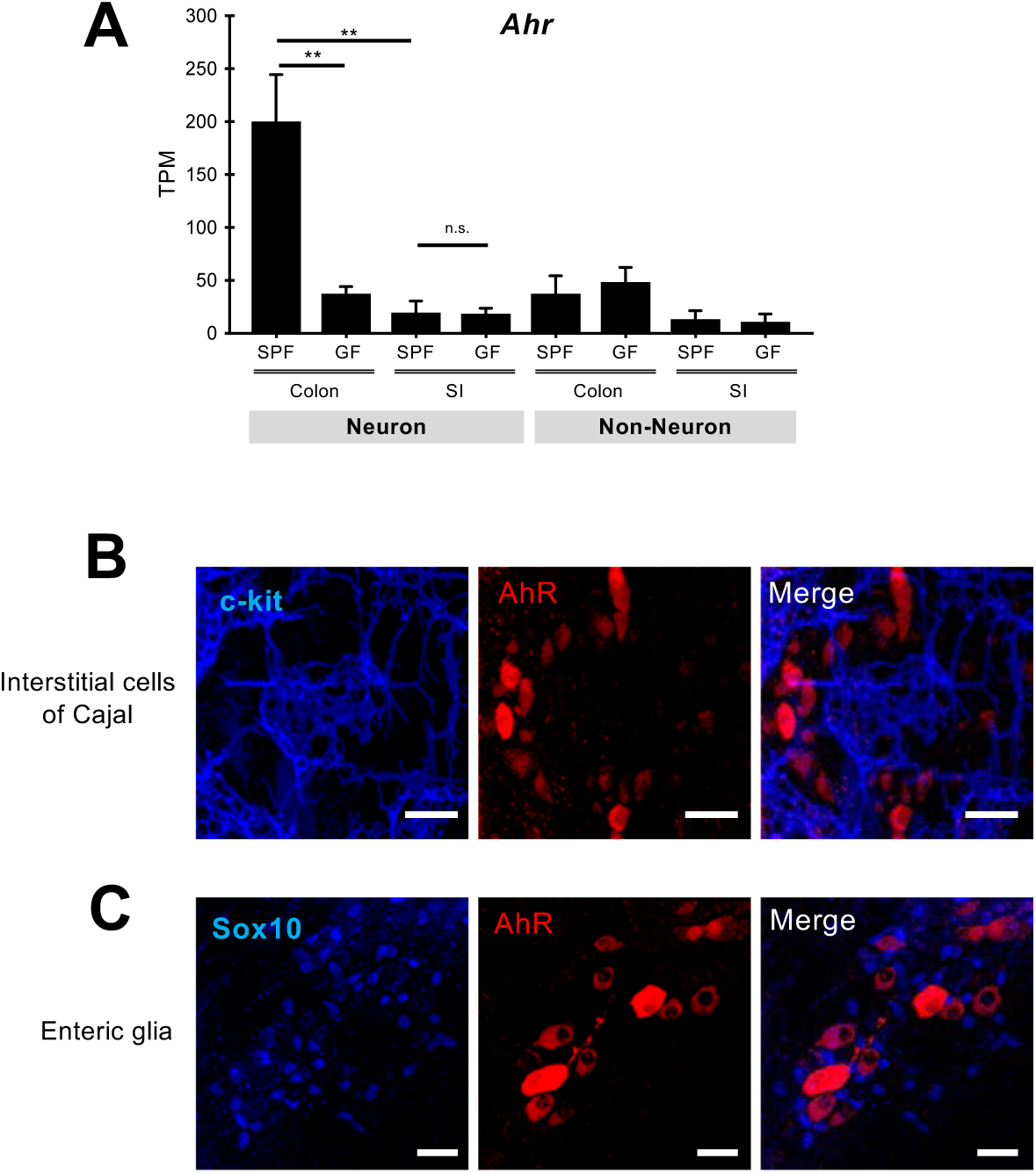
In muscularis externa *AhR* is expressed predominantly by neurons. (**A**) Quantification of AhR transcripts (TPM mean ± SEM, n=4 SPF and 3 GF mice) by nRNAseq in neuronal and non-neuronal nuclear preparations from the muscularis externa of the colon and SI of SPF and GF mice. *p* values were determined by one-way ANOVA test followed by Tukey’s test. ** *p<0.01,* n.s: not significant. TPM, transcripts per kilobase million. (**B** and **C**) Representative images of colon myenteric ganglia immunostained for c-Kit (a marker of interstitial cells of Cajal) and AhR (B) and Sox10 (a marker for enteric glial cells) and AhR (C). Scale Bars: 30µm.

**Figure S6.**
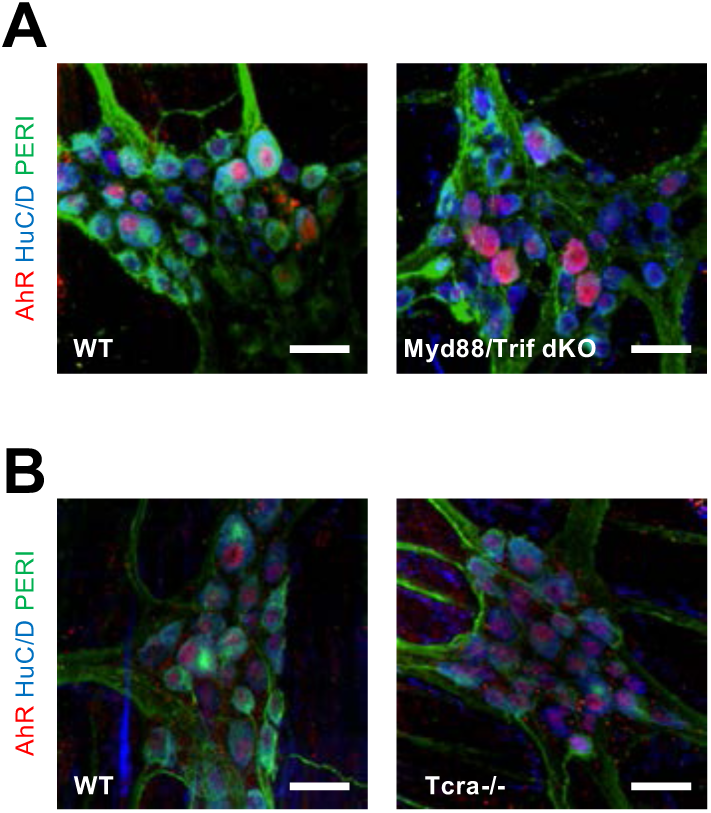
T cells and TLR signaling are dispensable for neuronal expression of *AhR*. (**A**) Immunostaining of colon myenteric ganglia from wild-type and *Myd88/Trif* double mutant GF mice inoculated with SPF microbiota (4 weeks prior to tissue harvesting) for AhR (red), HuC/D (blue) and PERI (green). (**B**) Immunostaining of colon myenteric ganglia from wild-type and *Tcrα*^*-/-*^ mutant (C57Bl/6) mice for AhR (red), HuC/D (blue) and PERI (green). Scale Bars: 30µm. PERI, peripherin.

**Figure S7.**
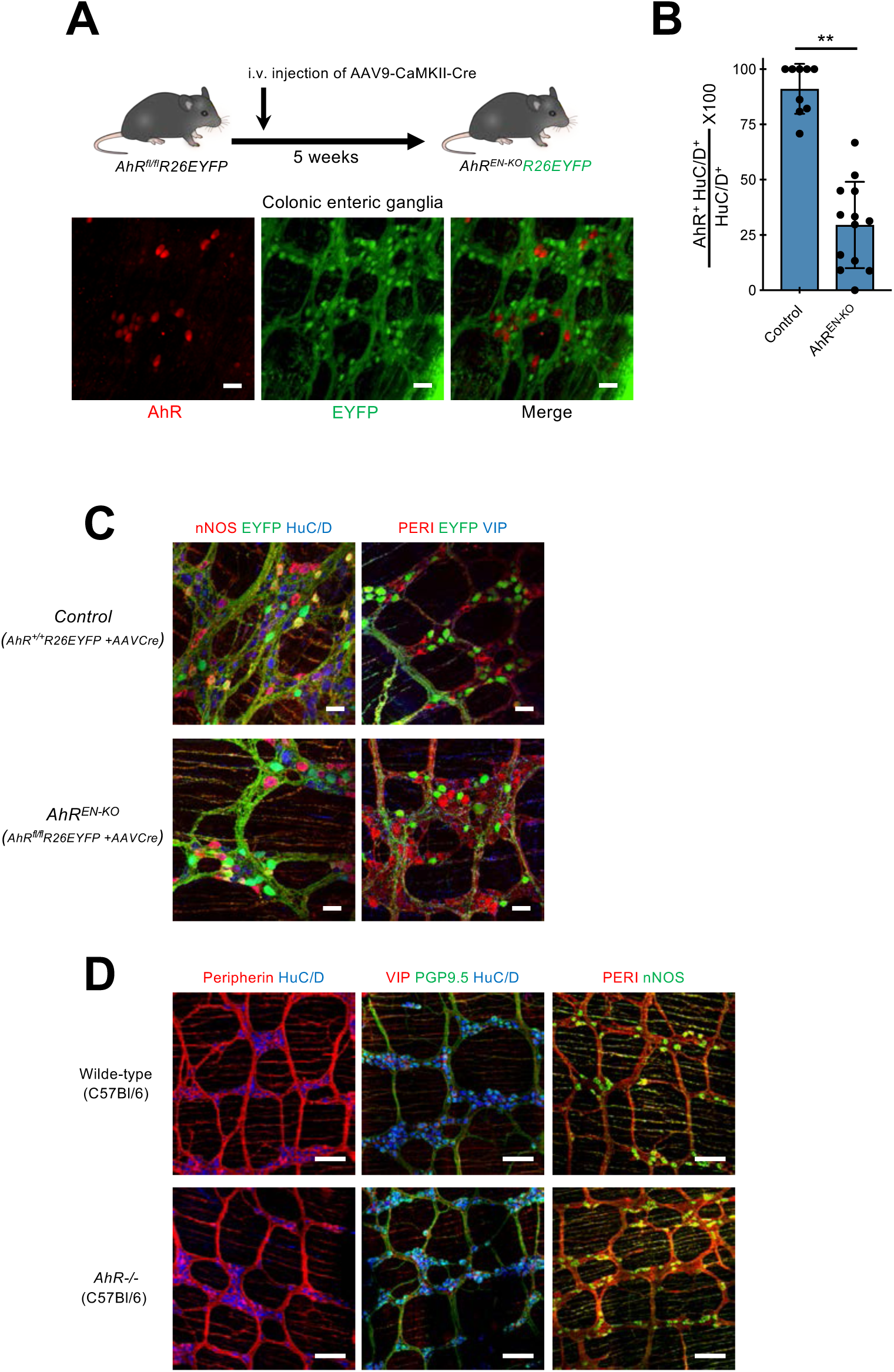
Deletion of *AhR* does not alter the organization and composition of myenteric ganglia. **(A)** Experimental design to delete *AhR* from enteric neurons of *AhR*^*fl/fl*^*;R26EYFP* mice. 4 weeks following intravenous injection of AAV9-CaMKII-Cre vector into *AhR*^*fl/fl*^*;R26EYFP* mice colon muscularis externa preparations were immunostained for AhR (red) and GFP (green). Note the lack of overlap between green and red cells. Data are representative of two independent experiments. Scale Bars: 30µm. (**B**) Group data (mean ± SEM) showing that administration of AAV9-CaMKII-Cre vectors to *AhR*^*fl/fl*^*;R26EYFP* mice results in dramatic reduction in the fraction of AhR^+^ enteric neurons in comparison to *AhR*^*+/+*^*;R26EYFP* mice injected with the same vector (control). Random images were acquired from the colon of each biological replicate (n=9 for control, n=13 for AhR^EN-KO^), and the average percentage of AhR^+^ HuC/D^+^ cells in total HuC/D^+^ neurons was calculated. Statistical test is Student’s *t* test. ** *p<0.01*. (**C**) Immunostaining of muscularis externa preparations from the colon of control (top) and AhR^EN-KO^ (bottom) mice with nNOS, EYFP and HuC/D (left) or PERI, EYFP and VIP (right). Scale bars: 30µm (**D**) Immunostaining of muscularis externa preparations from the colon of wild-type (top) and AhR^−/−^ (bottom) mice with PERI and HuC/D (left), VIP, PGP9.5 and HuC/D (middle) and PERI and NOS (right). Scale bars: 100µm. PERI, peripherin.

**Figure S8.**
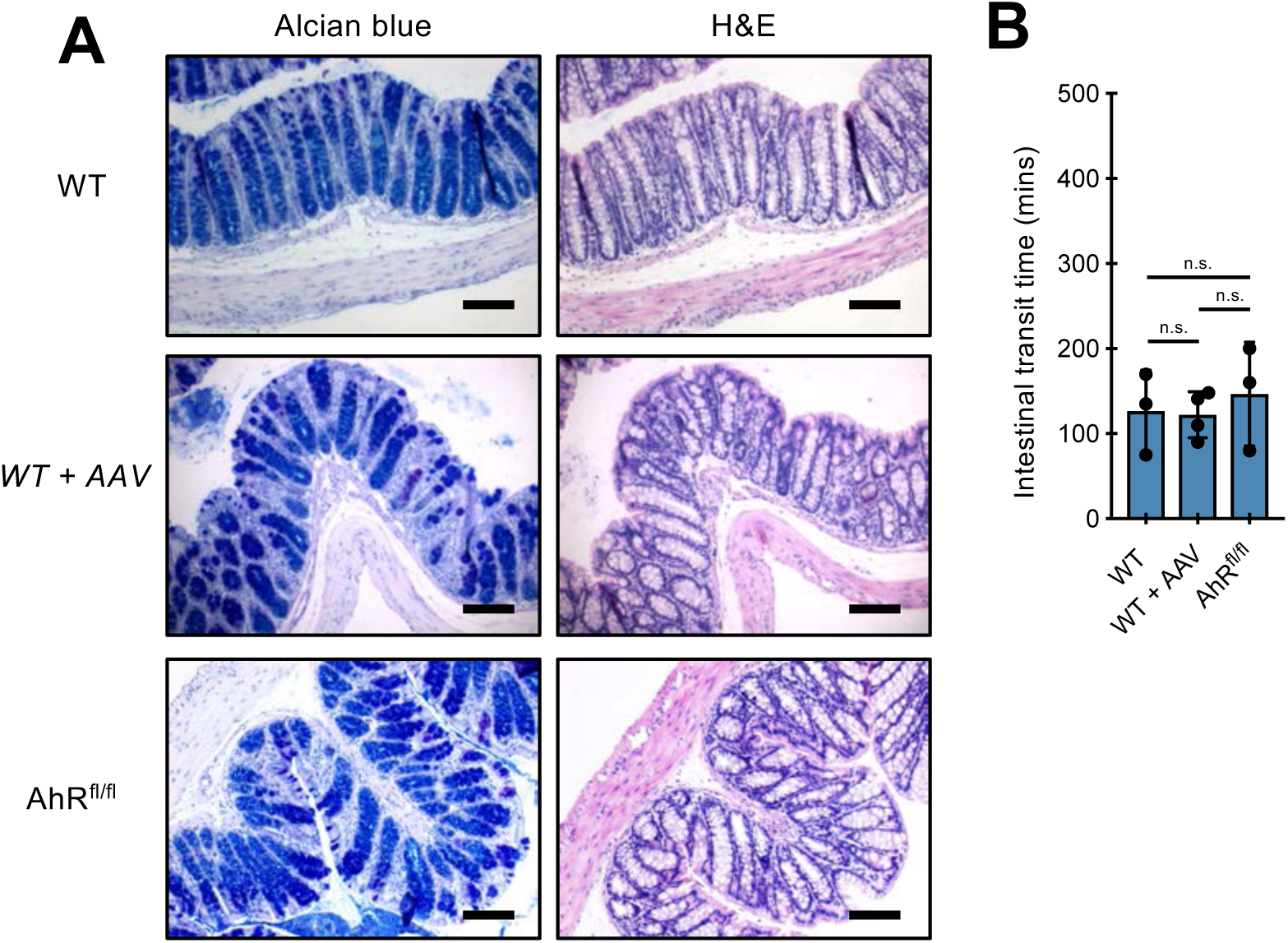
Systemic administration of AAV vectors to mice does not elicit an inflammatory response or intestinal dysmotility. (**A**) Cross sections from the colon of wild-type (top), wild-type infected with AAV9-CaMKII-Cre (middle) and *AhR*^*fl/fl*^ (bottom) mice stained with Alcian blue-PAS (left) or hematoxylin-eosin (H&E) (right). (**B**) Group data (mean ± SEM) showing that AAV-CaMKII-Cre vector injection into wild-type mice is not sufficient to alter intestinal transit time. n=3 (WT), 4 (*WT+AAV*), 3 (AhR^fl/fl^). Statistical test is one-way ANOVA test followed by Tukey’s test. n.s: not significant.

**Figure S9.**
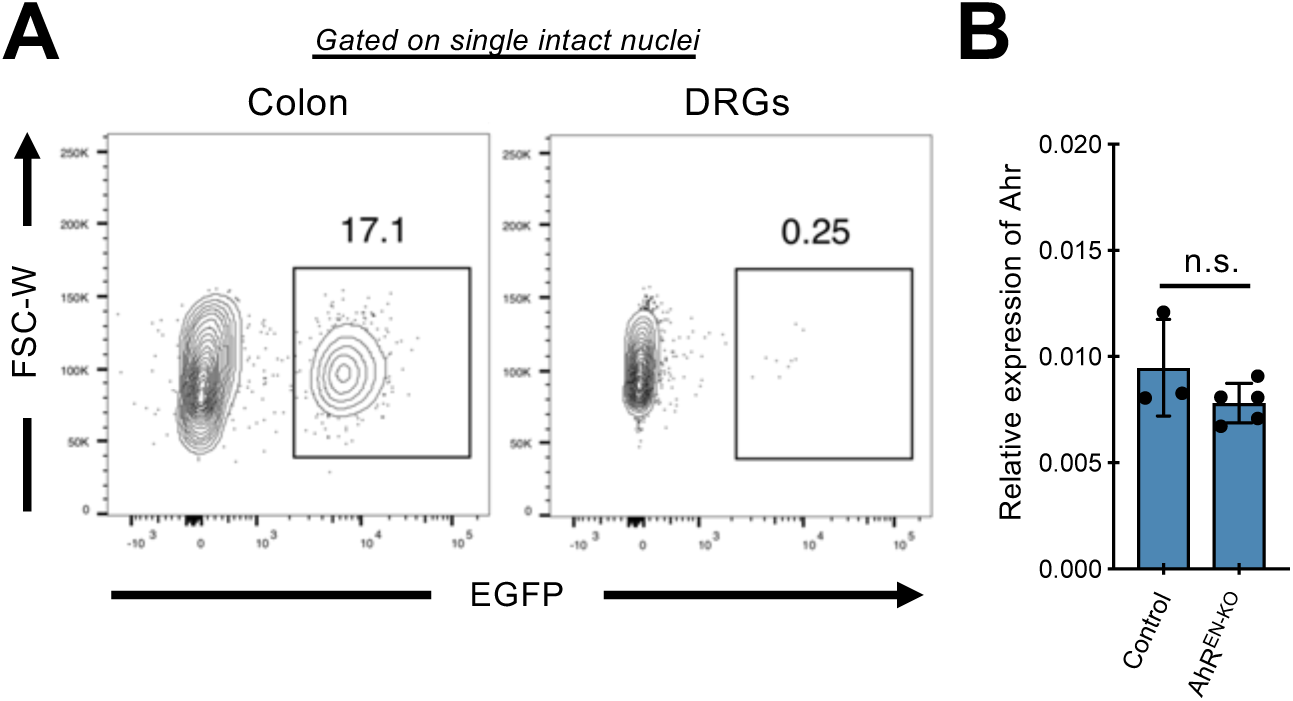
Minimal targeting of DRG neurons by intravenous administration of AAV vectors. (**A**) Representative FACS plots of nuclei from colon muscularis externa (left) and thoracic and lumber DRGs (right) of C57BL/6 mice injected with AAV9-CaMKII-EGFPKASH 5 weeks prior to nuclear preparation. Plots were gated on single intact DAPI^+^ nuclei. (**B**) Group data (mean ± SEM) showing similar expression of *Ahr* in thoracic and lumber DRGs from control and *AhR*^*EN-KO*^ mice. n= 3 (control) and 5 (*AhR*^*EN-KO*^). Statistical test is Student’s *t* test. n.s: not significant.

**Figure S10.**
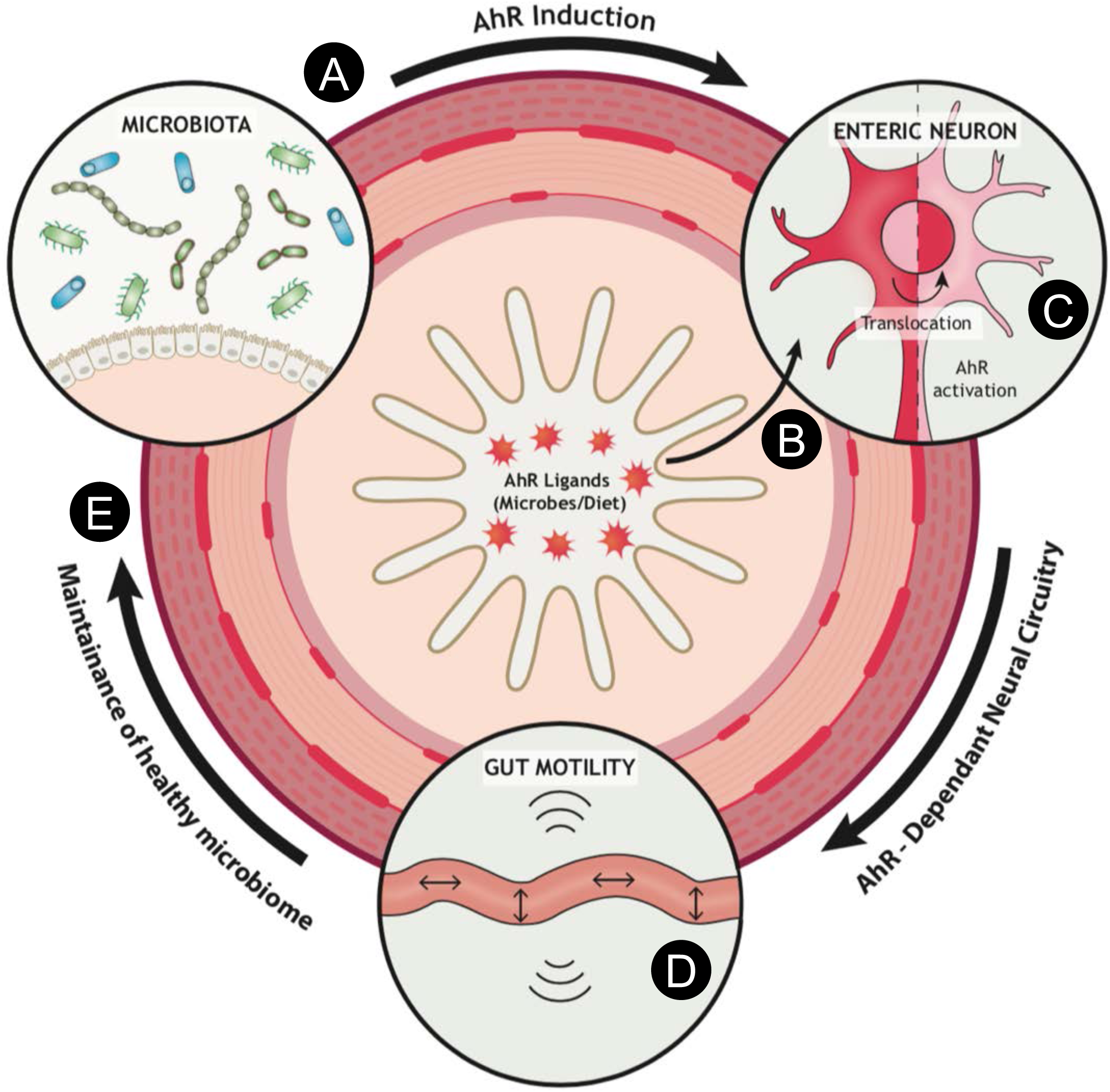
Model depicting the contribution of neuron-specific AhR activity in maintaining a healthy microbiome. Microbiota colonization upregulates AhR in the majority of enteric neurons in the distal intestine (**A**). This enables neural networks to monitor the lumen or tissue environment of the gut for the presence of metabolites (derived for example from microbiota or diet) that function as natural AhR ligands (**B**). Upon ligand binding, cytosolic AhR translocates into the nucleus inducing AhR-dependent transcriptional programs (**C**) that in turn modulate intestinal motility (**D**). Normal peristaltic activity of the gut is crucial for maintaining a healthy microbiota composition and avoiding abnormal bacterial overgrowth and dysbiosis that can be detrimental to gut homeostasis and host defence (**E**).

**Table S3.**

| Primary antibodies |  |  |  |  |  |
| --- | --- | --- | --- | --- | --- |
| Antibody | Company | Catalog no. | Host | Clonality | Dilution |
| α-AhR | Enzo Life Sciences | BML-SA550-0100 | Rabbit | PC | 1:200 |
| α-cKit | R&D Systems | AF1356 | Goat | PC | 1:200 |
| α-Calbindin | Abcam | ab82812 | Mouse | MC | 1:200 |
| α-Calretinin | Swant | CG1 | Goat | PC | 1:200 |
| α-ChAT | Millipore (Chemicon) | AB144P | Goat | PC | 1:200 |
| α-EGFP | Abcam | ab13970 | Chicken | PC | 1:200 |
| α-HuC/D | Invitrogen | A-21271 | Mouse | MC | 1:200 |
| α-mCherry | Abcam | ab167453 | Rabbit | PC | 1:200 |
| α-NF-M | Proteintech | 66396-1-Ig | Mouse | MC | 1:200 |
| α-nNOS | Abcam | ab1376 | Goat | PC | 1:200 |
| α-Peripherin | Santa Cruz | sc-7604 | Goat | PC | 1:200 |
| α-PGP9.5 | Biogenesis | 7863-0504 | Rabbit | PC | 1:1000 |
| α-S100β | DAKO | z-0311 | Rabbit | PC | 1:500 |
| α-Sox10 | Santa-Cruz Biotechnology | sc-17342 | Goat | PC | 1:200 |
| α-Tuj1 | Biolegend | MMS-435P (801201) | Mouse | MC | 1:200 |
| α-VIP | Abcam | ab22736 | Rabbit | PC | 1:200 |

| Secondary antibodies |  |  |  |  |  |
| --- | --- | --- | --- | --- | --- |
| Antibody | Company | Catalog no. | Host | Clonality | Dilution |
| α-Mouse Alexa Fluor 647 | Invitrogen (Life Technologies) | A-31571 | Donkey | PC | 1:500 |
| α-Rabbit Alexa Fluor 647 | Invitrogen (Life Technologies) | A-31573 | Donkey | PC | 1:500 |
| α-Goat Alexa Fluor 568 | Invitrogen (Life Technologies) | A-11057 | Donkey | PC | 1:500 |
| α-Goat Alexa Fluor 647 | Invitrogen (Life Technologies) | A-21447 | Donkey | PC | 1:500 |
| α-Goat Alexa Fluor 488 | Invitrogen (Life Technologies) | A-11055 | Donkey | PC | 1:500 |

**Description of RNAscope® (Advanced Cell Diagnostics) probes**
| **Probe** | **Channel** | **Catalog no.** |
| --- | --- | --- |
| *Ret* | C1 | 431791 |
| *Ahr* | C1 | 452091 |
| *Pou3f3* | C1 | 441521 |
| *Pde1c* | C1 | 489011 |
| *Ano5* | C1 | 557141 |
| *Pantr2* | C1 | 483721 |
| *Unc5d* | C2 | 480461 |
| *Col25a1* | C3 | 538511 |

